# miRge3.0: a comprehensive microRNA and tRF sequencing analysis pipeline

**DOI:** 10.1101/2021.01.18.427129

**Authors:** Arun H. Patil, Marc K. Halushka

## Abstract

MicroRNAs and tRFs are classes of small non-coding RNAs, known for their roles in translational regulation of genes. Advances in next-generation sequencing (NGS) have enabled high-throughput small RNA-seq studies, which require robust alignment pipelines. Our laboratory previously developed miRge and miRge2.0, as flexible tools to process sequencing data for annotation of miRNAs and other small-RNA species and further predict novel miRNAs using a support vector machine approach. Although, miRge2.0 is a leading analysis tool in terms of speed with unique quantifying and annotation features, it has a few limitations. We present miRge3.0 which provides additional features along with compatibility to newer versions of Cutadapt and Python. The revisions of the tool include the ability to process Unique Molecular Identifiers (UMIs) to account for PCR duplicates while quantifying miRNAs in the datasets and an accurate GFF3 formatted isomiR tool. miRge3.0 also has speed improvements benchmarked to miRge2.0, Chimira and sRNAbench. Finally, miRge3.0 output integrates into other packages for a streamlined analysis process and provides a cross-platform Graphical User Interface (GUI). In conclusion miRge3.0 is our 3^rd^ generation small RNA-seq aligner with improvements in speed, versatility, and functionality over earlier iterations.

## INTRODUCTION

MicroRNAs (miRNAs) are a group of small non-coding RNA that act as master regulators of gene translation (1). Altered miRNA expression affects cell stress signaling, cell proliferation, and is central to human disease states (2). Each miRNA can target scores of mRNAs and can regulate entire signaling pathways (3). In humans, the number of miRNAs is controversial with only 1,111 miRNAs reported in miRGeneDB 2.0 (4), but 2,656 miRNAs in the miRBase database (version 22) (5). Other groups list hundreds or thousands of additional small RNAs which are non-canonical miRNAs (6–8). The exploration of miRNA expression profiles has added valuable insights into mechanisms of cell processes in health and disease (2,9).

tRNA fragments and halves (tRFs) have more recently become of interest through their role in pathophysiology (10). They are thought to increase based on cell stressors and have variable expression patterns based on cell type and method of cell stress (11). Some functions overlap with miRNA activities. Other interactions with RNA binding proteins impact on diseases such as cancer (12).

Advances in genomics has enabled cost-effective high-throughput sequencing from small RNA libraries to study tissue (13,14) and cell (8,15) expression. Within small RNA-seq datasets, in addition to miRNAs and TRFs, other types of RNA such as rRNA, siRNA, snoRNA, and mRNA fragments exist, some of which are variable in disease (16). In order to accurately identify and quantify sequence data, the reads must be aligned to reference databases with appropriate parameters. Additionally, for miRNAs, a collection of nearly similar sequences, termed isomiRs, sum up to the reads of a particular miRNA (17).

In 2015, our laboratory published a small RNA-seq alignment tool, miRge, focused on miRNA expression (18). It was designed as a fast, smart alignment tool coded in Perl. An update, miRge2.0, coded in Python 2.7, introduced new features including novel miRNA discovery, isomiR description in the GFF3 format, tRNA fragment characterization and detection of A-to-I changes (17,19). With the deprecation of Python 2, this could not be maintained. Here we present miRge3.0, coded in Python 3, as an even faster, more robust tool with greater functionality, unique molecular identifier (UMI handling) and a new graphical user interface (GUI) for both input and output data.

## MATERIAL AND METHODS

The revised small-RNA analysis pipeline miRge3.0 is implemented in Python (v3.8) and is designed to run in Linux, macOS, and Windows 10 with windows subsystems for Linux (WSL). miRge3.0 can be called from a command line or run through a GUI described below. miRge3.0 uses Cutadapt (v3.0) (20) for adapter trimming on both ends of FASTQ reads and downstream quality control. Pandas (v0.25.3) libraries have been implemented to enable memory efficient annotations of small RNA molecules by reducing I/O-based operations. miRge3.0 requires Bowtie (v1.3.0) (21), ViennaRNA (v2.4.16) (22), SAMtools (v1.7) (23), biopython (v1.78), sklearn (v0.23.1), numPy (v1.18.4), SciPy (v1.4.1) and reportlab (v3.5.42) for alignment operations, novel miRNA discovery, and GUI visualizations. In addition to this, miRge3.0 is designed to handle reads with Unique Molecular Identifiers (UMI) and further integrates DESeq2 (release 3.12) (24) function in R (v4.0) to compute differential expression. The entire package is available through Bioconda (25), Python package index (PyPi) repository and the source code is available at GitHub (https://github.com/mhalushka/miRge3.0). The package is also available as a docker image at https://quay.io/ (26).

### Workflow and Alignment Steps of miRge3.0

The initial step of miRge3.0 is the removal of bad quality reads and adapter sequences using Cutadapt. miRge3.0 allows for a wide range of adapters (both 5’ and 3’ types) to be natively called and removed. At this step, UMIs, if present, can be trimmed for a fixed length specified by the user. Removal of PCR duplicates can be removed when the unique counts of UMI and read sequence combinations are recorded. After this initial step, identical reads are collapsed into a single read and the counts are captured in a Pandas data frame. The Pandas data frame will join reads and corresponding read counts for two or more FASTQ samples to create a complete data frame. This data frame is used for downstream alignments and for generating various summary results.

The multiple alignment steps, using Bowtie, of these small RNA reads is the same as described previously (18,19). There is an initial tight alignment to a modified miRNA library with no mismatches followed by alignments to libraries of hairpin miRNA, small nucleolar RNA (snoRNA), ribosomal RNA (rRNA) and other non-coding RNA allowing a single mismatch. Only the top best alignment for primary and mature tRNAs with 1 and 0 mismatches respectively are retained. The remaining unaligned reads are further aligned to messenger RNA (mRNA) library with 0 mismatches followed by a loose alignment to the original modified miRNA library allowing up to 2 mismatches to identify isomiRs. Bowtie is run across all libraries with ‘--norc’ option to restrict reads from alignment against the reverse-complement reference strand. miRge3.0 incorporates the latest new standard format for reporting isomiRs in GFF format (17). After each step, the data frame gets a status alignment tag of “1” where reads are aligned and “0” as not aligned. At each iteration, the collapsed reads get a tag of “1” and finally, two data frames with “1 – mapped” and “0 – unmapped” reads are created. Where, mapped data frame has annotations of small RNA (as columns) for which reads are aligned. This mapped data frame is used for A-to-I editing, isomiR analysis, tRF annotations, and generating Binary Alignment Map (BAM) files for visualization in the Integrative Genomics Viewer (IGV). The unmapped data frame is used for novel miRNA prediction. The overview of the workflow is shown in Figure 1.

**Figure 1:**
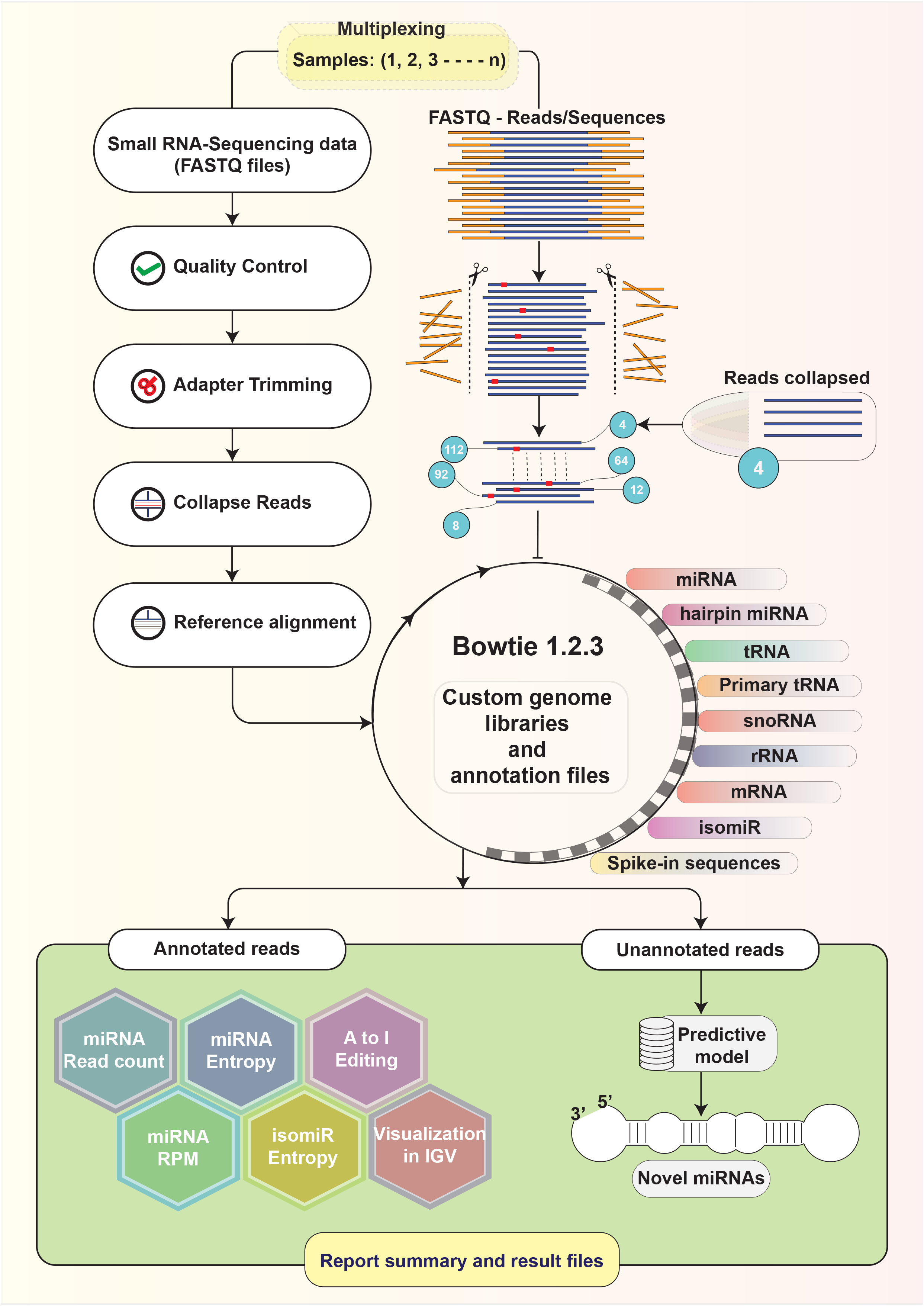
An overview of miRge3.0 workflow

### Novel miRNA prediction

Novel miRNA detection is based on a machine learning algorithm and on a prediction model, built using support vector machine (SVM). The features predicted in our previous version (19) were rebuilt using sklearn (v0.23.1) to support python 3. The model and novel miRNA prediction is as described previously (19). In brief, the unaligned reads are aligned and clustered to the genome. The most stable region of each cluster is extracted as a putative mature miRNA and the coordinates are set based on the cluster where the frequency of each base over the total number of reads in that cluster is greater than 0.8. Further, the pre-miRNA hairpins based on the fold energy of the sequence surrounding the cluster is determined using RNAfold. A SVM model is applied on the features of these pre-miRNAs and a probability value is calculated determining the significance of identified putative miRNA.

### miRge3.0 Graphical User Interface

The miRge3.0 suite offers a GUI with substantial ease to install and use. The cross-platform GUI is developed with Node package manager (NPM, v6.14.4), Node.js (v12.16.3) and Electron (v1.4.13). HTML forms the framework behind the Electron shell and CSS adds to the overall presentation of the framework, while the dynamic functionality of the tool is built with JavaScript. The output includes an HTML page that describes the summary of the data.

For running miRge3.0, the GUI menu allows all features of the command line request (Figure 2). This is done with the HTML tags to request user input parameters and these parameters are further assembled to a variable as a command line argument to execute miRge3.0. This variable is written as a shell script in Linux and Mac prior to calling miRge3.0, while, in Windows10, the miRge3.0 command along with all the parameters are run directly on the windows command prompt. A “progress” tab automatically appears showing the progress of miRge3.0 run as a function of JavaScript. An additional JavaScript function hides this tab and opens a “visualize” tab upon completion of the run describing the summary and results. All features related to switching between parameters is enabled by JavaScripts and HTML tags.

**Figure 2:**
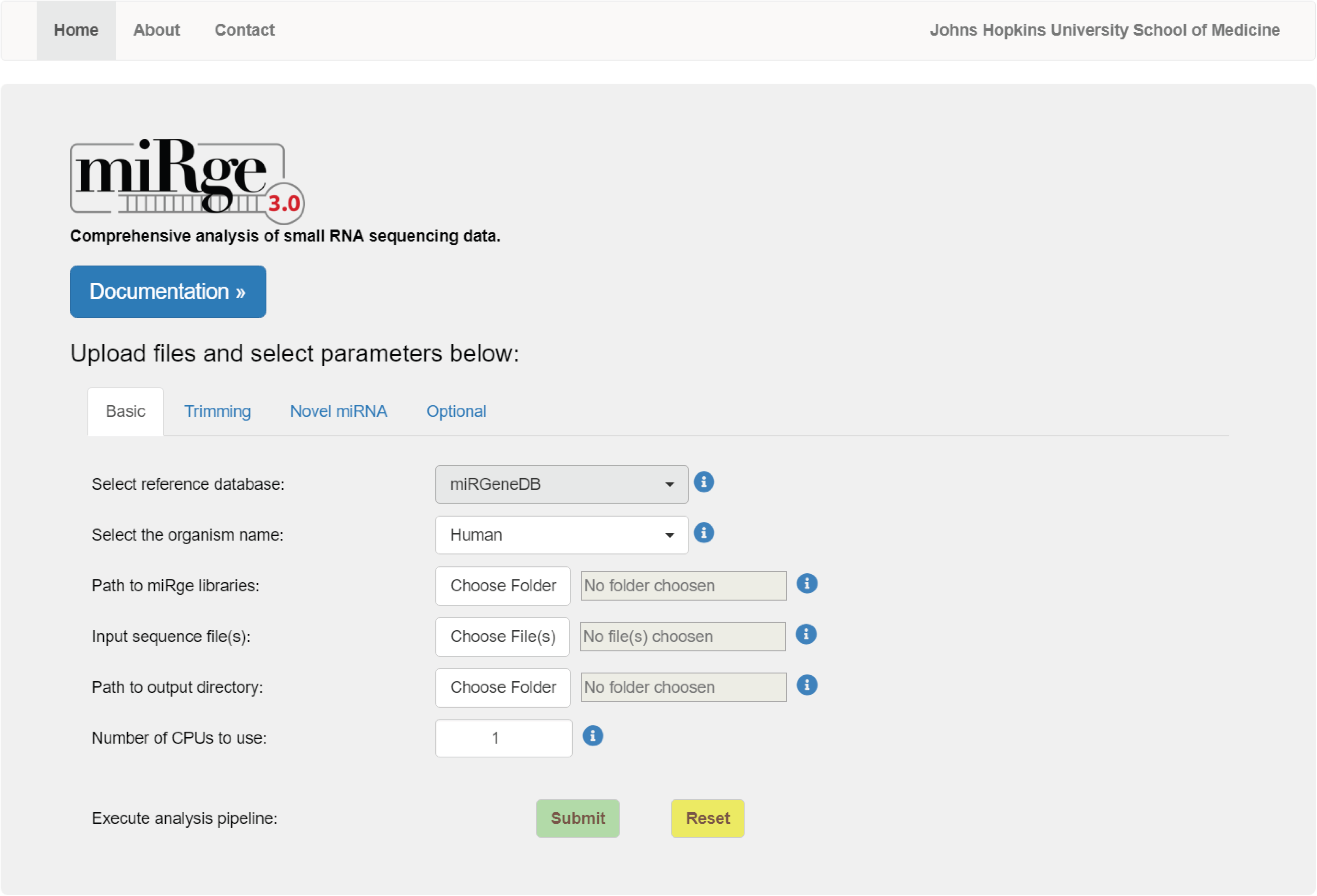
Screenshot of miRge3.0 – GUI

After the miRge3.0 run is complete, a dynamic summary output is generated (Figure 3). This includes tabs containing: 1) a distribution of small RNA as a stacked horizontal bar chart for each sample, 2) read length distributions as a histogram, 3) a donut chart representing the variant types and heatmap showing variant counts across the canonical sequence for the top 20 abundant miRNAs, 4) a tile map showing the top 40 abundant miRNA expressions, 5) a distribution of UMI counts as a histogram, and 6) a list of novel miRNAs identified. The interactive charts are rendered using JavaScript and CSS obtained from High Charts (https://www.highcharts.com/), the icons in the miRge visualization HTML tabs are obtained from Font Awesome (v5.8.2, https://fontawesome.com/) and the JavaScript for interactive HTML table depicting novel miRNAs is obtained from Data Tables (https://datatables.net/). This interactive HTML page is also available for Command Line Interface () users, and also has an option to save the results in various formats such as JPEG, SVG, PNG, PDF, and as tabular formats. The software tool is available for download and use through SourceForge (https://sourceforge.net/projects/mirge3/).

**Figure 3:**
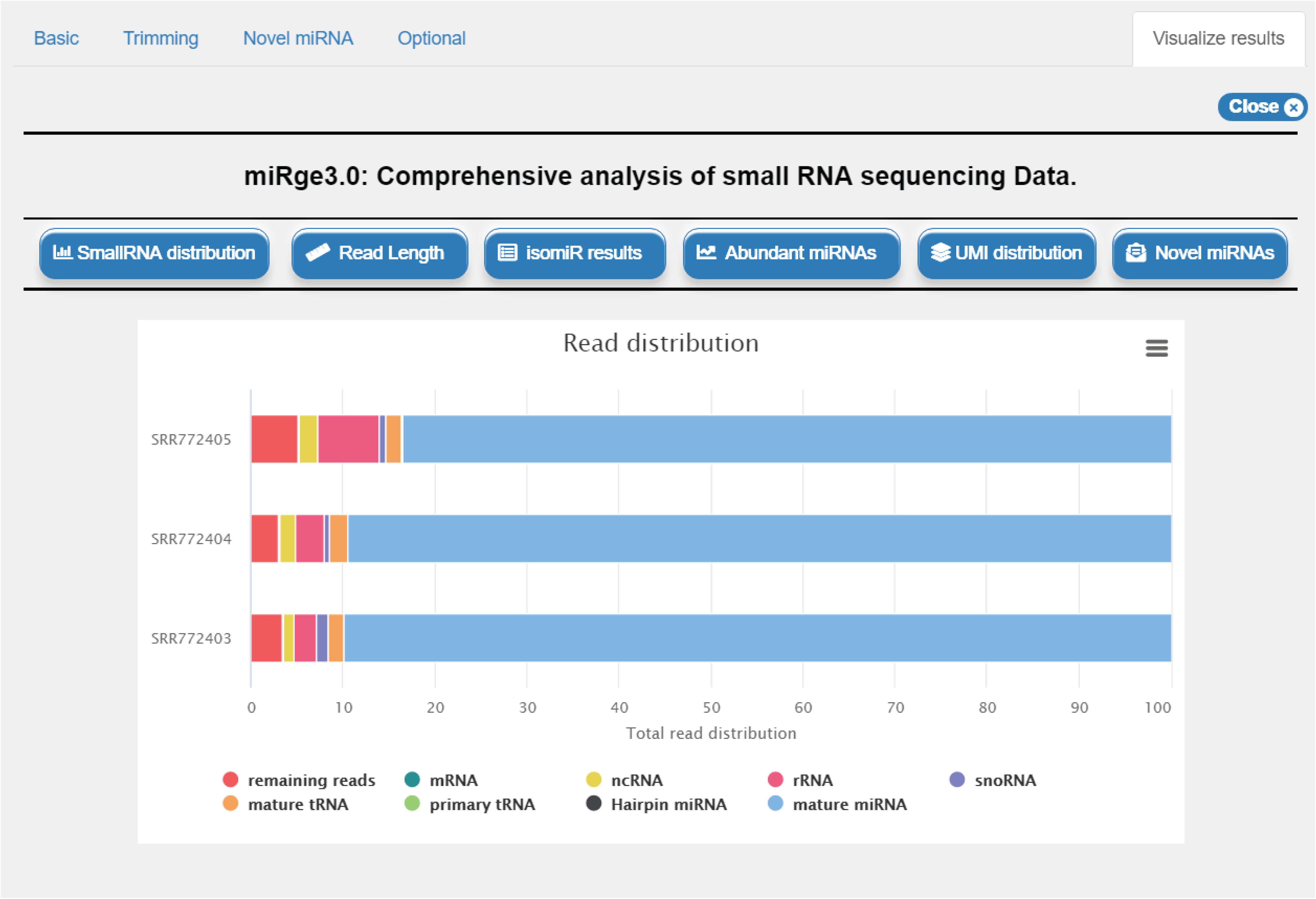
Visualization offered by miRge3.0

### Speed Testing

To benchmark miRge3.0 to other tools, we utilized the following publicly-available SRA files; ERR747967 (platelets) (27), SRR5127204 (cortical neuron), SRR5127209 (renal proximal epithelium), SRR5127210 (retinal pigment epithelium), SRR5127236 (cardiac fibroblasts) (8), SRR1853808 (CD8+ Neonatal T Cells) (28) and SRR1028924 (Islet Alpha Cells) (29). The miRge3.0 pipeline uses standard small RNA libraries as a reference and, to compare the performance, two tools with similar features, Chimira (30) and sRNAbench (31), were chosen. Table 1 describes some common technical features among the chosen tools. Speed comparisons were performed in two stages. At the first stage, the samples were run with default parameters for all tools except for specifying Illumina 3’-adapter sequence TGGAATTCTCGGGTGCCAAGGAACTCCAG. The samples were uploaded in FASTQ.gz format to Chimira and accession IDs were provided to sRNAbench to download from NCBI SRA and perform the annotation. The samples with FASTQ extension were run for miRge2.0 and miRge3.0. The annotations were timed from start to completion for all tools excluding the sample download and upload time. Chimira does not report time for Reaper and Tally steps in their log file, therefore, the timing for each sample were monitored manually. sRNAbench converts ‘FASTQ’ files to ‘FASTQ.gz’ using command ‘gzip’ before performing annotations. We have reported two time series including gzip conversion time (sRNAbench) and without (sRNAbench - gzip).

**Table 1:** Comparison of common technical features across tools

| **Features** | **miRge3.0** | **miRge2.0** | **Chimira** | **sRNAbench** |
| --- | --- | --- | --- | --- |
| Reference database | Modified libraries | Modified libraries | Hairpin | Genome or libraries |
| Input format | FASTQ, FASTQ.gz | FASTQ, FASTQ.gz | FASTQ.gz, FASTA.gz | FASTQ.gz, FASTQ, FASTA, read count, SRA |
| Batch processing of input files | Yes, enables to read files from folder | Yes | Yes (<2.6 Gb) | Yes |
| Detect novel miRNAs | Yes | Yes | No | Yes |
| Standalone CLI | Yes | Yes | No | Yes |
| Standalone GUI | Yes | No | No | Yes |
| Alignment tool | Bowtie | Bowtie | BLASTn | Bowtie |
| tRNA fragment annotation | Yes | Yes | No | No |
| BAM for visualization | Yes | No | No | No |
| Process UMI | Yes | No | No | Yes |
| Interactive graphics/visual output | both | Visual output | Interactive graphics | both |
| Image formats | JPEG, PDF, PNG and SVG vector image | PDF | Not available for download | PNG |
| isomiRs in GFF format | Yes | Yes | No | No |
| Differential expression | Yes | No | No | Yes |

Chimira and sRNAbench report modifications and isomiRs respectively along with basic annotations. In the second stage, although, the miRge2.0 and miRge3.0 pipeline reports isomiR counts for each miRNA, for consistency, the miRge2.0 and miRge3.0 pipelines were supplemented with the ‘-gff’ parameter to generate an isomiR GFF file. Further, Chimira and sRNAbench are hosted online and, while server specifications are unknown, they are expected to have large RAM capacity and numerous cores. sRNAbench uses 10 CPUs to download data from NCBI SRA and 4 CPUs for small RNA annotations. Thus, the miRge pipelines were run with 4 and 12 CPUs for each sample. Chimira has a maximum file upload limit of 1.6GB precluding the running of sample SRR1028924.

### Processing Unique Molecular Identifiers (UMIs)

The UMI for small RNA sequencing largely falls under two category, Illumina/NEXTFLEX/Clontech 4N method and Qiagen UMI sequencing. The methods vary in UMI location within the adapter sequences. The UMIs are processed in parallel during quality and adapter removal process, where a miRge3.0 function stores UMI sequence for each read until the downstream pre-processing is completed. These reads with UMIs for Illumina and Qiagen are handled differently. The parameter for handling 4N ligation adaptor/UMIs is ‘-umi x,y’ where x is the count of UMI nucleotides 5’ to the template and y is the count of nucleotides 3’ to the template (Figure 4A). The Qiagen method has two adapter sequences, and a UMI in between them (Figure 4B). The parameter ‘–qiagenumi’ identifies that the data is from Qiagen platform and distinguishes this from other platforms. The ‘–umiDedup’ parameter, if specified, removes duplicates of the identical UMI + Read combinations before collapsing with identical read counts.

**Figure 4:**
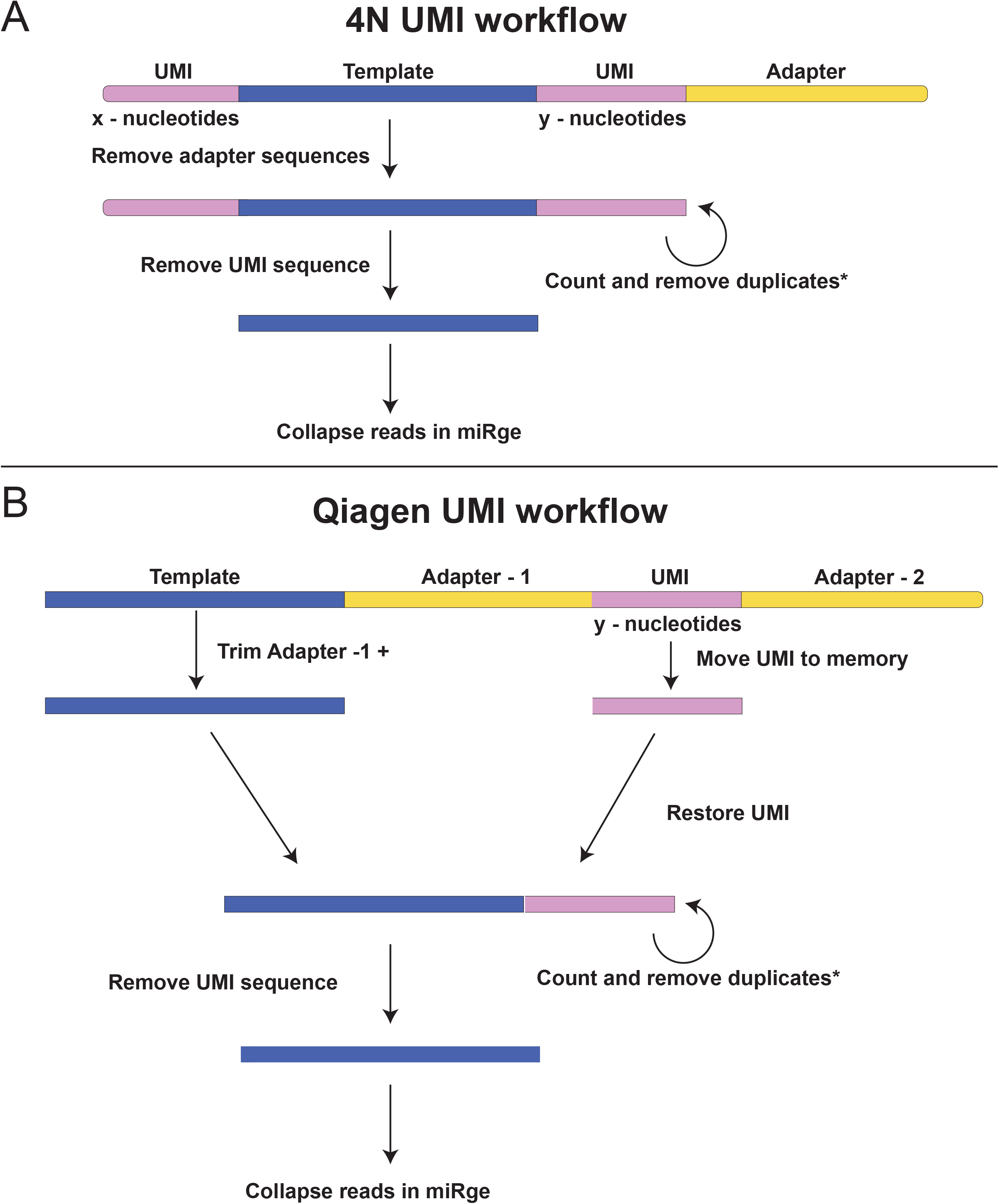
Workflow of processing UMIs. A) 4N ligation adaptors (UMI) on both sides of the template are collected and counted. *= if called by –umi, duplicates will be removed. B) Qiagen UMIs, within the adapters are collected and counted. *= if called by –umi, duplicates will be removed.

### Comparing 4N and Qiagen UMI output with or without deduplication

Correcting for counts of the 4N ligation adaptor/UMI was performed on SRR6379839 (synthetic construct) (32), and SRR9115360 (NEXTflex, Human Brain) (33) and a Qiagen processed sample, SRR8557389 (34). The parameters for 4N method include -a TGGAATTCTCGGGTGCCAAGG (for adapter removal), -umi 4,4 (to trim 4 bases on both ends of the read after adapter removal). The parameters for Qiagen UMI include -a AACTGTAGGCACCATCAAT (for adapter removal), --umiqiagen (specifying Qiagen), and -umi 0,12 (To trim 0 bases at 5’ and 12 bases at 3’ after adapter removal). While rest of the default parameters are kept constant for both runs, each dataset was run with and without the --umiDedup (remove duplicates) option.

### Differential expression analysis

Bioconductor R package DESeq2 (24), is integrated in miRge3.0 to estimate differential expression among the samples. A metadata of groups for control and condition should be supplemented in a file with tab-delimited text format, and miRNA read counts are used to perform the differential expression. If differential expression analysis is called, the analysis results appear in a text file reporting miRNA’s and its corresponding log2 fold change values along with the p-value and adjusted p-value. Further, PDF files reporting a volcano plot of differentially expressed miRNAs and principal components analysis (PCA) plot of the input samples are automatically generated. An RData file is also generated for further integration with other tools and/or edit the output graphical format using R.

### Other datasets and resources used

As described previously, a combination of JavaScripts, CSS, and HTML makes a dynamic interactive visualization chart. We have used SRR772403, SRR772404, SRR772405, SRR772406 (35) and SRR649562 (36) datasets for functionality testing throughout the development of the miRge3.0 and to generate the supplementary figures (1–3). The coordinates for transcripts with intron exon boundaries were formatted using GffRead during the preparation of the reference libraries (37). A python package for building custom libraries is available at https://github.com/mhalushka/miRge3_build.

### Hardware

Small RNA-seq analysis for miRge2.0 and miRge3.0 were performed with Intel Xeon CPU E5-2690 v4 @ 2.60GHz, 256GB RAM Ubuntu x86_64 GNU/Linux server and a laptop Eluktronics MAG-15, with 12 CPUs (Intel Core i7-9750H processor at 2.6GHz) and 64 GB DDR4 RAM was used to develop miRge3.0.

## RESULTS

### Features of miRge3.0

miRge3.0 is an updated/improved version over our previous version miRge2.0, with various changes being incorporated, the most significant ones among them are coding in Python 3, functionality with newer Cutadapt packages, differential expression determination, processing UMIs, and a GUI offering interactive graphical input and output. miRge3.0 also has a significant speed advantage over miRge2.0 due to efficient coding involving Pandas data frame and multiprocessing from python class ProcessPoolExecutor. The command line version of miRge3.0 can be easily installed using pip and conda installation procedures.

### Output files of miRge3.0

Multiple output files are generated in miRge3.0. There are both a file of mapped reads with counts for each of the small RNA types across samples processed and a similar file of unmapped reads. Separate files log read counts and reads per million (RPMs) for each miRNA species (miRBase or MirGeneDB). A log file is reported, that includes the information of the executed command and parameters along with time stamp of that execution. Additional output files depend on user arguments/requirements. A Pandas data frame produces additional files such as isomiRs in GFF3 format, a BAM file for visualization in IGV, tRNA fragment annotations, A to I editing, and detection of novel miRNAs. In addition, the mapped reads could be saved as FASTA files for individual input samples. Figure 5 depicts the BAM tracks on IGV representing mapped canonical and isomiRs reads. At the end of the execution, miRge3.0 reports an HTML and a JavaScript file dynamically generated to report the summary of the analysis. By default, this summary includes an interactive graphic of percent read distribution for each sample as a horizontal stacked bar-graph (Supplementary Figure 1A), a histogram showing read length distribution (Supplementary Figure 1B) and a honeycomb (tile map) showing the top 40 abundant miRNAs (Supplementary Figure 2A). If specified, it will further include: 1) a table of novel miRNAs identified across samples (Supplementary Figure 2B), 2) cumulative isomiR variant type distribution in pie chart (Supplementary Figure 3A) and heat map showing read distribution of isomiRs for the top 20 abundant miRNAs (Supplementary Figure 3B) and 3) a histogram showing the distribution of UMIs for each sample (Figure 6A-C). All of the summary file host interactive graphics allow users to view the data in a table format and enable the downloading of high-resolution images for presentation and/or publications.

**Figure 5:**
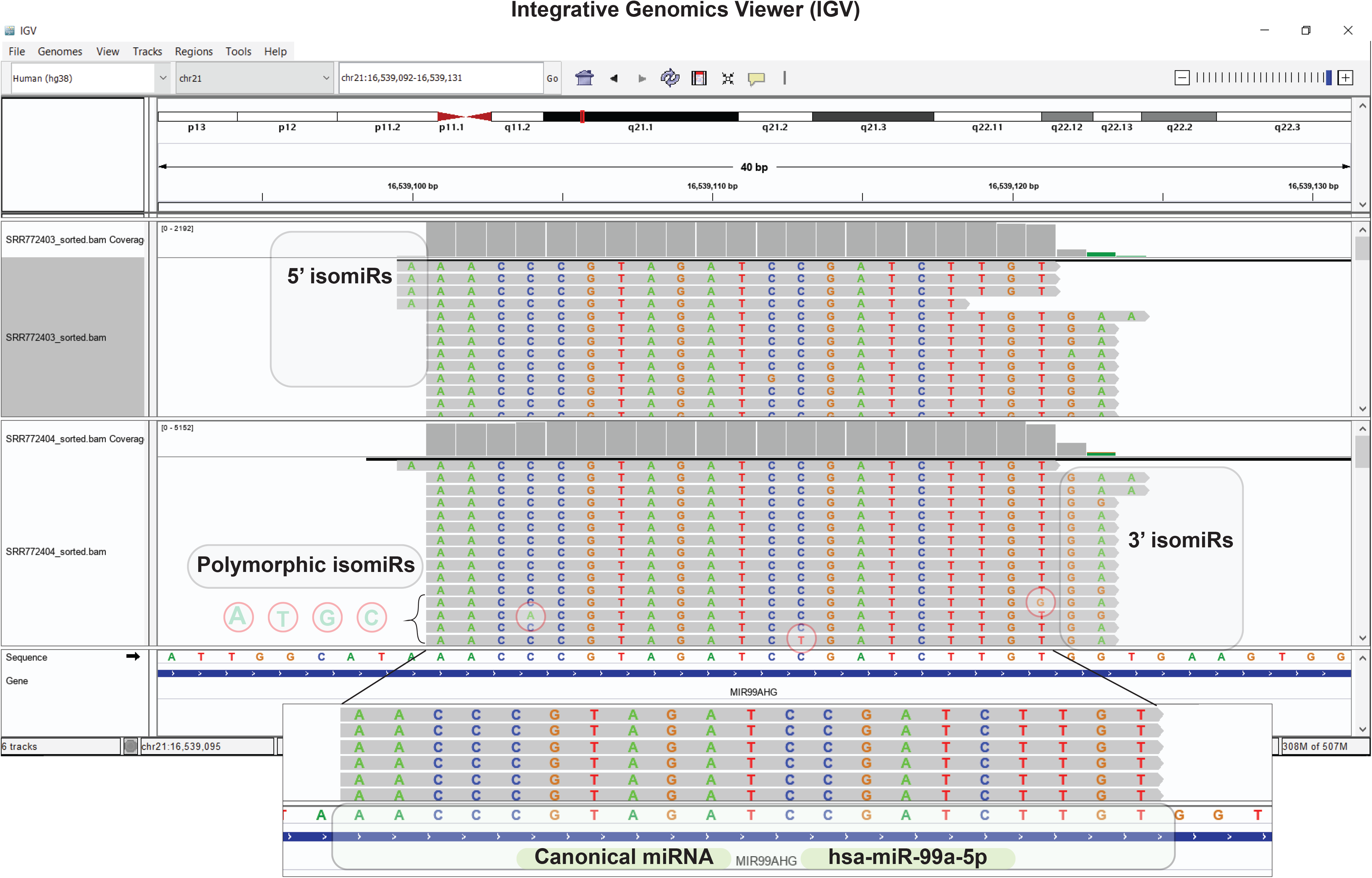
Screenshot of IGV showing canonical (miRNA) and variant (isomiR) reads

**Figure 6:**
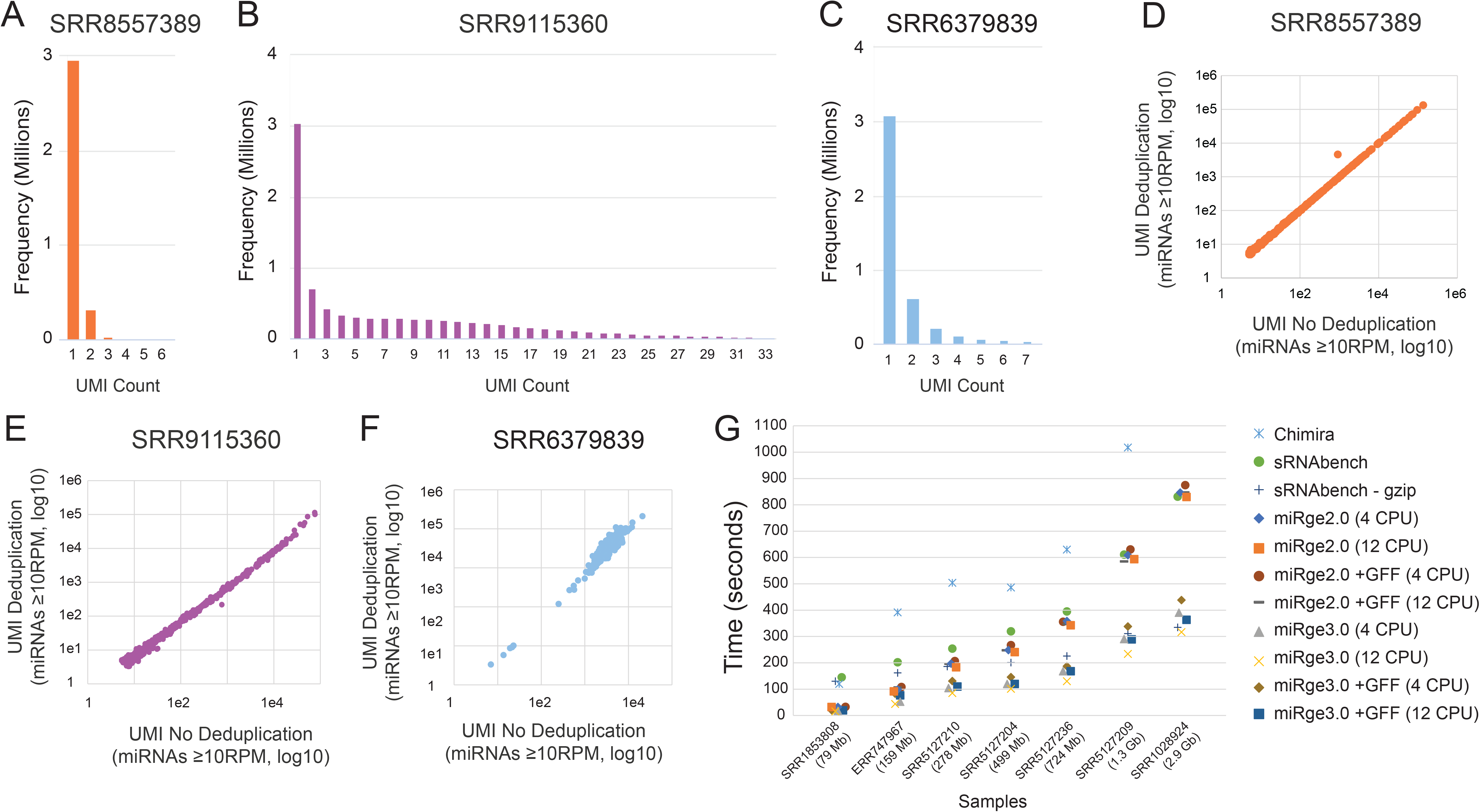
Comparison across tools and UMI analysis. A) A FASTQ file with Qiagen UMIs showing few examples of duplication. B) A FASTQ file with the 4N ligation adaptors showing significant counts of duplicated UMIs. C) A FASTQ file of synthetic data showing some increase in UMIs. E) The correlation between deduplicated and non-deduplicated miRNA RPM counts for Qiagen UMIs was very similar between repeated miRge3.0 runs (r^2^= 0.9996). D and F) The correlation was lower for samples with 4N UMIs (r^2^= 0.95 and 0.81, respectively). G) Run speed for 7 samples ranging from 79MB to 2.9GB file sizes across 4 tools.

### UMI deduplication analysis

miRge3.0 has the ability to detect, remove, and adjust for UMIs. Currently, two types of UMIs exist, those that act as ligation adaptors at the ends of the template (4N, Figure 4A) and those within the adapter (Qiagen, Figure 4B). The 12 bp Qiagen UMI, when collapsed with the entire read sequence, shows only rare duplications (Figure 6A). Due to ligation interactions of the shorter length 4N adaptor, the complete sequencing of short miRNA reads, along with the overall fewer nucleotides of the UMI (N=8), many more copies of these 4N-style ligation UMIs can be seen in real (Figure 6B), or synthetic (Figure 6C) data. It is more likely the “birthday paradox” is impacting on UMI distribution, rather than true amplification error (https://bitesizebio.com/35201/probability-theory-molecular-barcodes/). As a result of these differences, UMI removal, as performed for Qiagen-type longer UMIs has little effect on read counts (Figure 6D), whereas UMI removal on 4N samples can have a more significant effect (Figures 6E, F). Therefore, the adjustment for 4N UMIs, likely overcorrects and undercounts abundant miRNAs (33).

### Speed comparison to other alignment tools

Performance speed of miRge3.0 was compared with miRge2.0, Chimira and sRNAbench. SRR datasets with different read depths used to monitor processing speed across these tools. As sRNAbench converts a raw “FASTQ” file into a “FASTQ.gz” file in an initial step, we provide time data for both options (sRNAbench and sRNAbench – gzip). miRge2.0 and 3.0 times were reported using 4 CPU or 12 CPU with or without the +GFF feature. Although all of these tools are fast relative to most other miRNA aligners, overall, miRge3.0 with 12 CPUs consistently had the best execution speed (Figure 6G). Non-default parameters, such as isomiR GFF reporting will compromise miRge3.0 speed.

### Abundance estimation across tools

We ascertained the abundance of miRNAs from multiple files across the four alignment tools to understand how similar the findings were from each method. These results are expressed in terms of raw read counts or as normalized abundance (RPM). Table 2 shows the number of unique miRNAs expressed and those with ≥10 RPM for seven samples across the four alignment tools. sRNAbench, miRge2.0 and miRge3.0 all show similar trends in the reporting of miRNA counts. Chimira has much higher detection of miRNAs, which is partly the result of it not having the same false-positive controls employed by the other tools such as minimal read counts or percent canonical read controls.

**Table 2:** miRNA annotations across tools

| Tissue/Cell | SRA References | Alignment Tool | miRNA Reads | Unique miRNAs | miRNAs > 10 RPM |
| --- | --- | --- | --- | --- | --- |
| T Cell CD8+ (Neonatal) | SRR1853808 | Chimira | 1,011,733 | 620 | 250 |
|  |  | sRNAbench | 996,785 | 396 | 190 |
|  |  | miRge2.0 | 1,004,944 | 317 | 216 |
|  |  | miRge3.0 | 1,004,438 | 318 | 217 |
| Platelets | ERR747967 | Chimira | 2,644,772 | 970 | 280 |
|  |  | sRNAbench | 2,030,951 | 504 | 222 |
|  |  | miRge2.0 | 2,652,547 | 423 | 256 |
|  |  | miRge3.0 | 2,561,109 | 423 | 259 |
| Retinal pigment epithelium | SRR5127210 | Chimira | 5,951,602 | 1,045 | 427 |
|  |  | sRNAbench | 5,274,077 | 911 | 318 |
|  |  | miRge2.0 | 5,802,230 | 777 | 372 |
|  |  | miRge3.0 | 5,742,333 | 777 | 371 |
| Cortical neuron | SRR5127204 | Chimira | 15,447,871 | 1,433 | 325 |
|  |  | sRNAbench | 15,076,370 | 1,033 | 243 |
|  |  | miRge2.0 | 15,353,234 | 873 | 296 |
|  |  | miRge3.0 | 15,316,966 | 875 | 295 |
| Cardiac fibroblast | SRR5127236 | Chimira | 7,759,550 | 1,427 | 410 |
|  |  | sRNAbench | 7,489,281 | 989 | 320 |
|  |  | miRge2.0 | 7,674,927 | 844 | 367 |
|  |  | miRge3.0 | 7,661,528 | 849 | 367 |
| Renal proximal epithelium | SRR5127209 | Chimira | 35,894,433 | 1,830 | 395 |
|  |  | sRNAbench | 34,961,340 | 1,382 | 306 |
|  |  | miRge2.0 | 35,748,118 | 1,171 | 359 |
|  |  | miRge3.0 | 35,697,299 | 1,178 | 360 |
| Islet Alpha Cell | SRR1028924 | Chimira | -- | -- | -- |
|  |  | sRNAbench | 43,746,104 | 1,177 | 240 |
|  |  | miRge2.0 | 43,880,787 | 910 | 279 |
|  |  | miRge3.0 | 43,770,112 | 913 | 279 |

## DISCUSSION

miRge3.0 is a significant advance from our earlier, popular miRge and miRge2.0 alignment tools. The major enhancements are increased speed due to additional multithreading, the development of GUI assistance for both setting up the alignment parameters and for viewing the output, handling of UMIs, and usability on MacOS, Linux, or Windows 10 platforms. This is in addition to useful features that were already in miRge2.0 which include a tRF detecting tool, native utilization of the GFF3 isomiR format, novel miRNA detection, and A-to-I editing detection. As a result of these features, we are confident that miRge3.0 is among the most useful small RNA aligners currently available.

A key aspect of all versions of miRge has been the use of specialized RNA libraries and iterative alignments. Aligning miRNAs and tRFs is challenging due to isomiRs, short lengths, modifications, and multiple related sequences duplicated in the genomes (10,18). Tradeoffs have to be considered in setting up alignment parameters as no one method can accurately assign all sequences. We have found that a “one size fits all” alignment method to the species’ genome results in the poorest understanding of the sequenced material with the most error. Alignments are improved when reducing the search space by focusing on only the transcribed parts of the genome. The second improvement we take is to perform multiple sequential searches of the collapsed RNA sequences to specific RNA species in which the alignments start with very “tight” parameters and ultimately become “loose” to account for isomiRs.

Overall, miRge3.0 is a more user-friendly tool with advantages over our previous version and other tools. However, there are a few limitations. The 4N-style UMI analysis can benefit from a computational method to adjust UMI correction relative to expect number of UMIs based on the overall short lengths. This and other miRNA alignment tools could experience speed enhancements by incorporating GPUs instead of CPUs for computational steps. These improvements are planned in additional iterations of miRge3.0.

In conclusion, miRge3.0 is a significantly improved version of our miRNA alignment tool that we believe has the strongest complement of speed, usability, accuracy, and features in this software category.

## AVAILABILITY

The source code is available at https://github.com/mhalushka/miRge3.0

Conda package for miRge3.0 is available at https://anaconda.org/bioconda/mirge3

PyPi package for miRge3.0 is available at https://pypi.org/project/mirge3/

Docker image is available at https://quay.io/repository/biocontainers/mirge3

The cross-platform GUI is available at https://sourceforge.net/projects/mirge3/files/

## ACKNOWLEDGEMENTS

The authors thank the Pathology Department IT department for providing a Mac computer for the developing and testing of miRge3.0 GUI on Mac OS (OS-X) and the use of the Ares Linux server (Pathology Informatics). The authors thank Kate Halushka with graphic assistance.

## SUPPLEMENTARY DATA

**Supplementary Figure 1:**
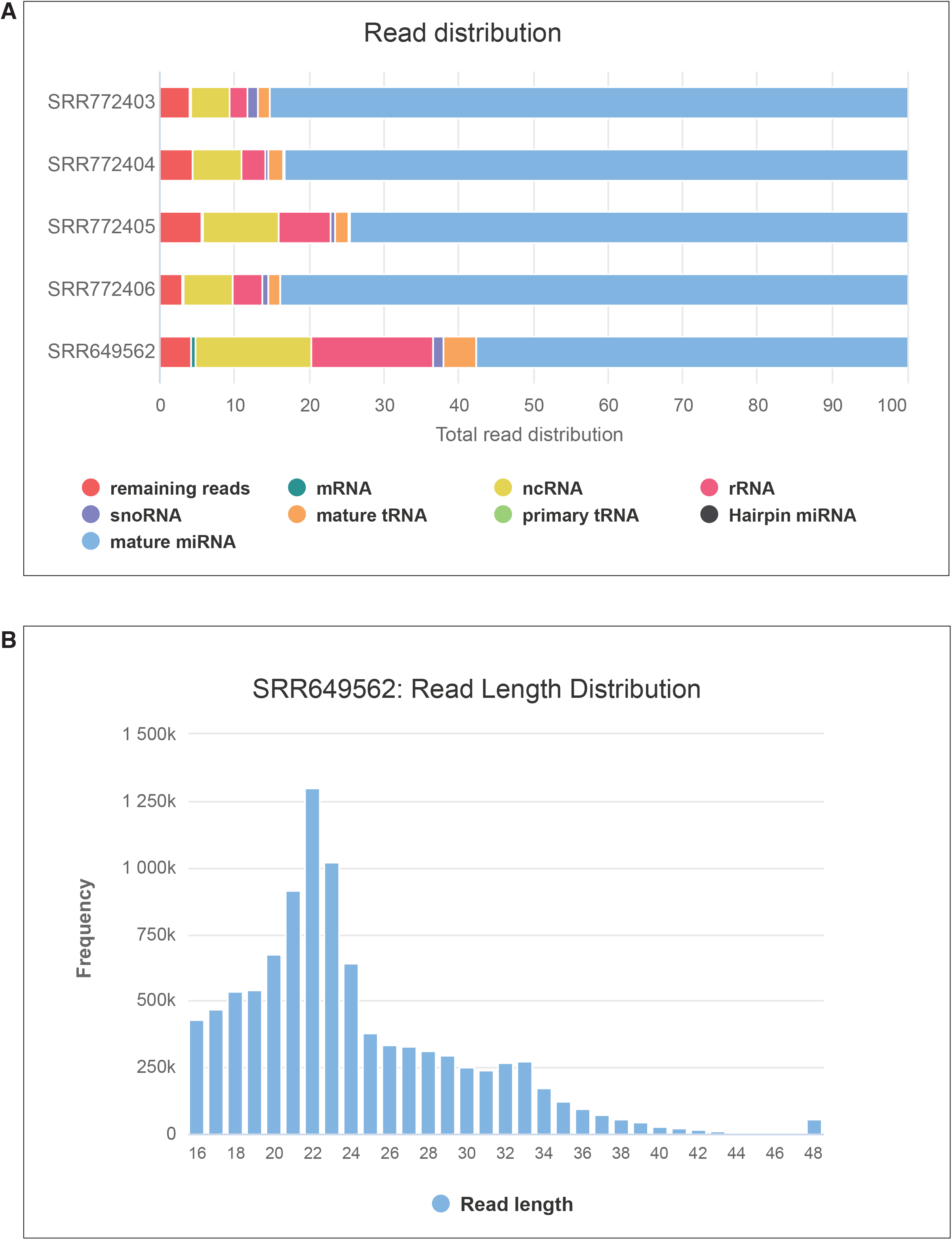
miRge3.0 visualization with minimalistic output showing annotation summary, A) total read distribution across small RNA types, B) Read length distribution

**Supplementary Figure 2:**
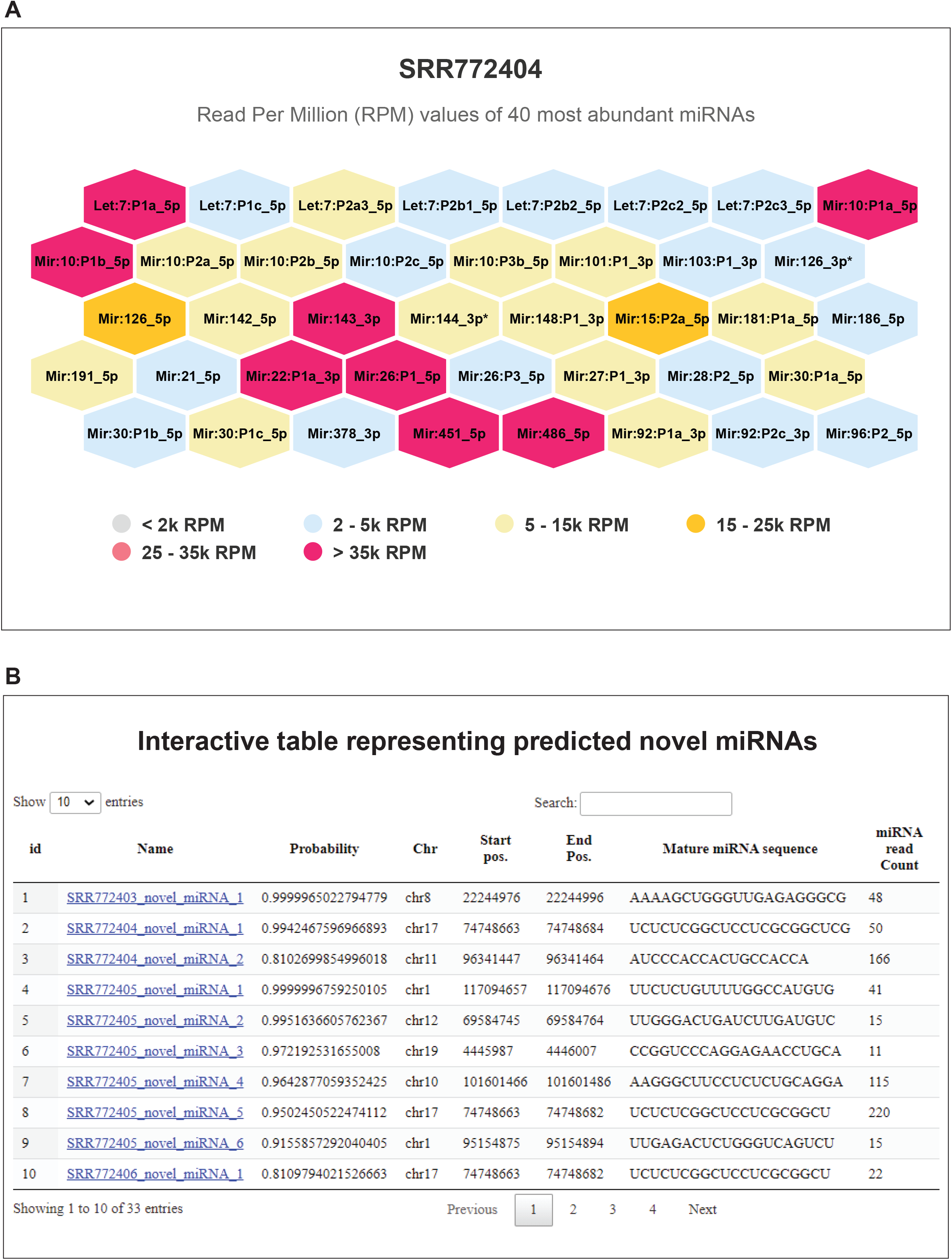
miRge3.0 visualization showing: A) Top 40 abundant miRNAs and B) table browser with predicted novel miRNAs

**Supplementary Figure 3:**
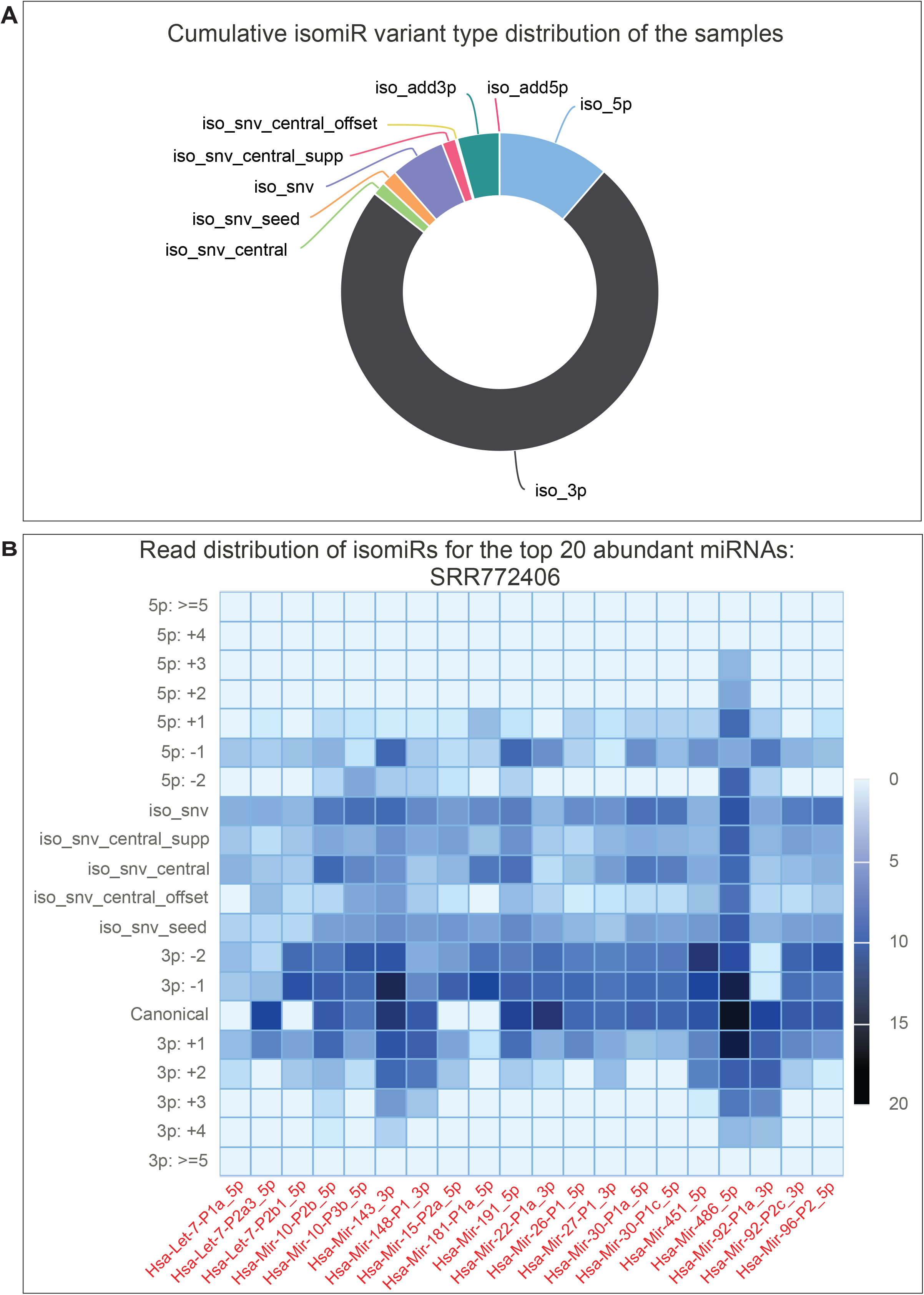
miRge3.0 visualization showing isomiRs as A) Cumulative distribution of variant types B) Heatmap showing read distribution of isomiRs for the top 20 abundant miRNAs

